# Residual Hearing Loss After Chronic Cochlear Implantation in Guinea Pigs is Primarily Conductive

**DOI:** 10.64898/2026.11.04.763992

**Authors:** Wenxuan He, Rubing Xing, Jordan D. Hill, Edward Porsov, Lina A. J. Reiss, George W. S. Burwood

## Abstract

It is desirous to preserve residual low-frequency hearing in cochlear implant recipients, as dual stimulation using electrical and acoustic signals results in superior speech-in-noise processing and musicality. However, cochlear implant recipients commonly lose their residual acoustic hearing gradually following surgical placement of the array. The governing mechanism is not understood and is likely due to multiple factors including surgical trauma, inflammation, and the foreign body induced immune response to the array, including formation of intracochlear fibrosis.

Normal mechanical function of the cochlea is expected to underpin residual hearing but has not been directly studied. This function includes efficient transmission of sound to the cochlea via the middle ear, sound propagation via the basilar membrane travelling wave, mechanical tuning characteristics, and the non-linear cochlear amplification of responses to low-frequency stimulation. Here, we examined these features in cochleae and middle ears of unimplanted, acutely and chronically implanted guinea pigs using our minimally invasive preparation and optical coherence tomography vibrometry.

We observed that mechanical tuning and amplification is not significantly altered by either acute or chronic implant. However, the response threshold was raised and proportional to middle ear responses. Additionally, the organ of Corti phase, and its sound intensity dependent changes, are unaltered by CI. Finally, a cluster analysis revealed sensitivity loss in chronically implanted animals is caused by middle ear dysfunction. Hence, in the guinea pig model, the dominant etiology of residual hearing loss is conductive. More research is required to establish whether residual hearing loss in humans similarly shares conductive elements.

## Introduction

Cochlear implants (CIs) are a highly effective means to restore hearing to patients with sensorineural hearing loss by bypassing the mechanoelectrical transduction apparatus, the organ of Corti, with direct stimulation of the spiral ganglion via a tonotopically stimulated array of electrodes (1). In patients where some low-frequency (residual) hearing is present (2, 3), electroacoustic stimulation can lead to more benefit compared to electrical hearing alone, with improvements to musicality and speech-in-noise intelligibility (4).

However, a subset of CI patients with residual hearing go on to progressively lose that hearing in the months to years following surgery (5, 6). Myriad factors such as acute inflammation, surgical trauma and foreign body response may contribute to this outcome (7–11), but as the cochlea is a mechanoelectrical transducer of sound, the involvement of cochlear mechanical changes remains a prominent potential component of poor residual hearing outcomes. Hitherto, modelling has suggested that direct damping of the basilar membrane by foreign body, be it fibrosis or CI array, can disrupt the hydromechanical function of the cochlea, and therefore lead to residual hearing loss (12). Additionally, modelling predicts a stiffening of the round window membrane also leads to low-frequency residual hearing loss (13). This modelling is corroborated by a lack of evidence that CI presence typically causes apical hair cell loss (14–16) outside of cases where exceptional insertion trauma (e.g. spiral ligament damage (17)) is apparent. In addition, local stiffness changes have been found at the level of the basilar membrane using atomic force microscopy (18).

Observing intact apical hair cells in guinea pigs, coupled with the modelling derived evidence for conductive, rather than sensory elements of residual hearing loss, lead to the hypothesis that chronic CI will cause conductive hearing loss but cochlear amplification will remain intact at the site of residual hearing, the cochlear apex. The mechanical function of a cochlear apex with a CI electrode present has not been investigated directly, and indirect measurements of organ of Corti function, such as auditory brainstem response (ABR) are less reliable for low frequencies in rodents, making the physiological state of the cochlear apex difficult to discern (19). Placement of the CI does not appear to impact low-frequency (below 1 kHz) neural responses, while inducing reversible conductive hearing loss for higher frequencies (20). Chronic CI can dilapidate distortion product otoacoustic emissions (DPOAEs) in mice, while the condition of the apical turn OHCs was not reported histologically (21). Under these conditions, it is not possible to separate a conductive deficit from a sensorineural one. It is vital to measure the mechanical properties of the middle ear and the apical organ of Corti directly, in order to achieve that separation.

Hence, we directly measured the sound evoked mechanical function of the umbo and apical turn organ of Corti in implanted guinea pigs, a popular CI model system (14, 15, 22–24). We report optical coherence tomography vibrometry (OCTV) measurements taken before and immediately after CI placement in one cohort of guinea pigs, and 8-10 weeks following CI in a second cohort. These measurements represent the first data of its kind.

We measured from the cellular component of the organ of Corti approximately 20% of the distance from the helicotrema, using our minimally invasive mirror preparation (25) and denoising technique. At sound pressure levels close to the hearing threshold, these measurements capture the cycle-by-cycle contribution of the outer hair cells to amplification of the base-to-apex travelling wave set up by pressure differences across the scalae in response to acoustically evoked middle ear motion (26). At higher sound pressure levels, the outer hair cell contribution is saturated, and the measured displacement is dominated by passive mechanics (27–29). The phase behavior of this travelling wave, largely underpinned by the passive mechanical properties of the cochlear partition, is also extracted. Previous studies in guinea pigs have demonstrated that the mechanical tuning of the cellular organ of Corti is comparable to that of the basilar membrane while the latter exhibits less amplification (30, 31). Additionally, the high frequency shoulder of the mechanical filter function is dominated by passive basilar membrane mechanics (27, 29).

We observe that, in the acute group, the apical organ of Corti itself was not strongly impacted by placement of the implant. However, some changes in the function of the middle ear were detected. In the chronically implanted group, we found that the apical organ of Corti gain was present, but middle ear function was often compromised, leading to a significant conductive component to the residual hearing loss. In both acute and chronic groups, there was no evidence of CI induced changes in organ of Corti tuning or non-linear phase behavior, even after separating out likely residual hearing loss subjects. We conclude that conductive hearing loss is a dominant component in our guinea pig model of residual hearing loss, representing a global rather than local change in cochlear hydromechanics and energy propagation efficiency. Further research is needed to identify the source of this conductive deficit, and to explore whether this also occurs in humans.

## Results

### Mechanical characteristics of the cochlear apex and middle ear before and after acute implantation

Contemporary surgical goals include hearing preservation for patients with residual hearing. In order to assess the impact of CI alone upon the function of the cochlear apex, and to validate our CI surgery in terms of hearing preservation, we made acute CI surgeries on anesthetized guinea pigs and recorded the mechanical response from the cochlear apex and from the umbo of the tympanic membrane before and after CI via cochleostomy.

Figure 1 shows the sound evoked mechanical response of the apical organ of Corti and middle ear pre-CI (blue), post-CI (red), and postmortem (black) for four animals given acute CI. To facilitate CI placement, the OCT system had to be moved after pre-CI measurements. Care was taken to move the OCT back to the original recording location, using visual alignment markers on the OCT microscope camera. The in-ear sound system was also re-calibrated following CI placement. Fig. 1A-D shows organ of Corti iso-level displacement tuning curves before and after CI, and postmortem (with CI in place).

**Figure 1:**
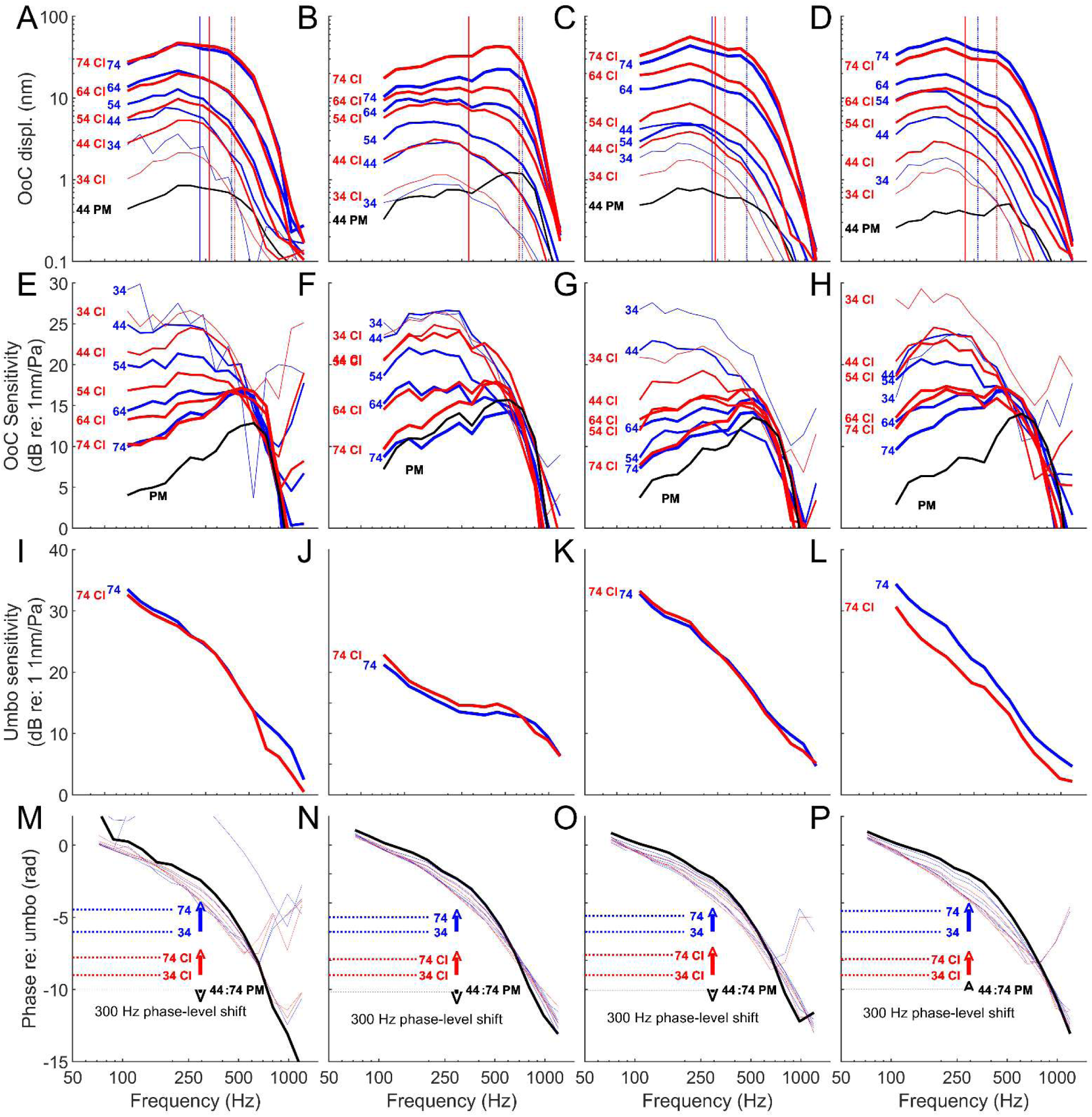
A-D: Raw iso-intensity displacement functions of the organ of Corti in 4 animals before (blue lines) and immediately after (red lines) placement of a CI, and after death (black lines). −3 dB re: peak displacement is indicated before (blue vertical lines) and after placement of the CI (red lines), for 44 dB SPL (solid lines) and 74 dB SPL (dashed lines). E-H: Iso-intensity Organ of Corti sensitivity functions (corrected for Umbo response) of the preparations in A-D before (blue lines) immediately after (red lines) CI placement, and postmortem in the presence of the CI (black lines). I-L: Iso-intensity sensitivity functions of the umbo from the same 4 preparations before (blue lines) and after (red lines) placement of the CI. M-P: Phase of each of the 4 preparations before (blue lines) and after (red lines) CI, and after euthanasia (black line). The level dependent phase difference between threshold and 74 dB SPL responses is shown using the dashed horizontal lines grouped using the same colors as the phase plots. The direction of that phase change is indicated by colored arrows.

Pre-CI measurements indicate band-pass tuning, the general shape of which did not change after CI. The following is a qualitative description of the impact of acute CI on the middle and inner ear mechanical response.

#### Raw displacement (Acute Cohort)

The amount of raw displacement evoked by the lowest decibel sound pressure level (dB SPL) stimulus (34 dB SPL) was reduced after CI in 3/4 animals (Fig. 1A, C, D). In one animal, the low SPL displacement increased slightly (Fig. 1B). In one case (Fig. 1 D), the absolute displacement at the highest SPL (74 dB SPL) was reduced after CI. In one case (Fig. 1A) the displacement remained stable, and in the remaining cases (Fig. 1 B, C) it increased.

#### -3 dB cutoff frequencies (Acute Cohort)

The local mechanical properties of the apical basilar membrane are unlikely to change upon acute implantation. To test this hypothesis, we calculated the −3 dB cutoff re: peak frequencies of the iso-intensity displacement turning curves at 34 dB SPL (Fig. 1, solid vertical lines) and 74 dB SPL (dotted vertical lines) before and after the CI was placed (blue and red respectively). The mean 34 dB SPL −3 dB cutoff re: peak was 280.62 ±60.73 Hz (N=4) before CI and 307.9±36.84 Hz (N=4) after CI. At 74 dB SPL, the cutoffs were 471.76±104.90 Hz (N=4) before CI and 451.81±104.63 Hz (N=4) after CI. The pre-CI values are comparable to those we have previously reported for lower and higher dB SPL stimuli (27, 31). Importantly, there was no statistical difference between the cutoff values measured prior to and post CI (34 dB SPL: p=0.415; 74 dB SPL: p=0.785, df=3, Student’s two-tailed t-test). This experiment indicates that the mechanical behavior of the apical organ of Corti is not influenced by acute placement of the CI in the basal turn.

#### Organ of Corti Sensitivity (Acute Cohort)

The condition of the apical cochlear amplifier can be estimated by converting the raw organ of Corti displacement to dB re: the pressure measured by the microphone in the ear canal. Note, that some lever gain is applied by the middle ear itself, and so even the passive organ of Corti will exhibit some gain. The gain evoked by a 34 dB SPL tone complex was reduced in only one case (Fig.1 G) but was comparable pre- to post CI for the other 3 animals. Importantly, sensitivity decreased postmortem, indicating all measured organ of Corti responses were amplified both pre-and post CI.

Compression (e.g. the difference in gain at the lowest vs. highest SPL) was impacted by implantation. In all cases, compression was reduced, either due to increased gain at high SPL (Fig 1. E-H), or this and a loss of gain at lower SPL (Fig. 1H). In the former case, this occurrence points to linearization and loss of dynamic range without a loss of gain (e.g. recruitment). In the latter case, a loss of gain and linearization are encountered.

#### Umbo response (Acute Cohort)

While there is evidence of sensory hearing loss akin to temporary threshold shift in the organ of Corti responses above, CI insertion could impact the umbo response. In 1/4 cases, Umbo sensitivity was diminished following CI placement and recalibration of the sound stimulus (Fig. 1 L). This suggests that the placement of the CI led to conductive effects, which would impact compression threshold but apparently not maximum gain at the cochlear apex – where gain is lost at the lowest SPL, this should be due to sensory deficit. Importantly, a loss of compression occurred in one of the cases where pre and post CI umbo responses were similar (Fig. 1 K), indicating that these changes were mainly sensory in nature.

In summary, acute placement of a CI in the basal turn of the guinea pig cochlea leads to varied and mixed hearing loss with differing contributions from the sensory and conductive components of that loss. However, the CI can be acutely placed while largely preserving apical non-linearity and gain – indicating that the guinea pig model of chronic cochlear implantation allows for examination of residual hearing mechanics.

#### Phase of the organ of Corti response (Acute Cohort)

The mechanical response of the organ of Corti can be understood in terms of its displacement, but also the phase of that displacement, which provides information on the propagation timing of the travelling wave from the umbo to the organ of Corti measurement site, and how this may change due to our various conditions and stimulus parameters.

A key feature of the phase of the travelling wave is its level dependent shift. In the basal turn, there is a progressive phase lag with increasing SPL (32). In the apical turn, using the same type of stimulus, we previously showed that the phase-SPL relationship is reversed, with decreasing group delay (and thus less phase accumulation or a relative phase lead) with changing SPL – an effect that disappears with loss of cochlear non-linearity (33, 34). Fig. 1 M-P show the phase responses for the acutely implanted animals before (blue curves) and after placement of the CI (red curves). A black curve indicates the phase of the postmortem response. The horizontal dashed lines and arrows indicate the magnitude and direction of the phase shift before and after CI, and postmortem (Fig. 1 M-P, blue, red and black). All animals show that the phase shift is positive and remains so before and after CI but vanishes postmortem. This indicates that the component of the organ of Corti phase that is dominated by amplification is unaffected by acute CI.

### Mechanical characteristics of the cochlear apex and middle ear after chronic implantation

#### Apical organ of Corti raw displacement (Chronic cohort)

Chronic CI placement foments the effects of foreign body immune response, which lead to inflammation and buildup of fibrosis close to the CI. We implanted several cohorts of guinea pigs for between 8 and 10 weeks, and after this latency period we assessed their apical mechanical function using OCTV as in the acute case. Figure 2 shows the iso-intensity displacement functions of the organ of Corti, recorded at a similar location to those in Fig. 1. The in vivo displacements at low SPL (Fig. 2, thin blue lines) are consistently above those of the measurements made following euthanasia of the animal (Fig. 2, thin black lines), demonstrating that the cochlear amplifier was active in the cochlear apex in chronically implanted animals – an outcome which is consistent with acute implantation (Fig. 1A-D). However, the threshold of the lowest detectable displacement differed between animals (e.g. Fig. 2A: 34 dB SPL, versus Fig. 2F, 74 dB SPL).

**Figure 2:**
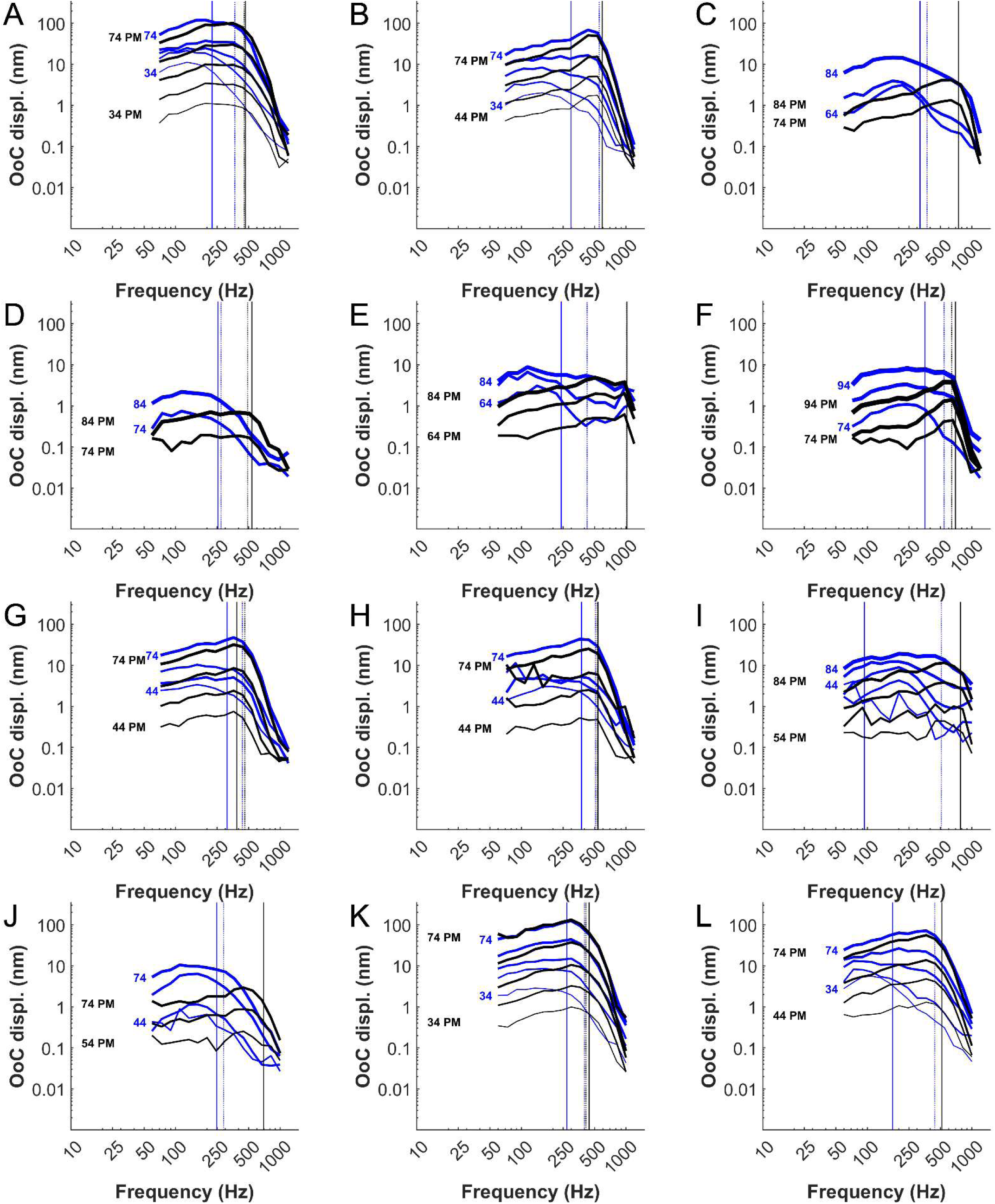
Raw iso-intensity displacement functions of the organ of Corti in 12 chronically implanted animals (A-L) before (blue lines) and at least 30 minutes after (black lines) euthanasia. −3 dB re: peak displacement is indicated before (blue vertical lines) and after euthanasia (black lines), for the lowest dB SPL (solid lines) and highest dB SPL (dashed lines) recorded.

#### -3 dB cutoff frequencies (Chronic Cohort)

Some evidence exists that supports the notion that chronic CI may induce stiffening of the guinea pig basilar membrane, even apical of the CI (18). This could potentially induce changes to the resonant properties and thus tuning of the apical organ of Corti. To test this, we estimated the −3dB cutoff re: peak displacement values for the chronically implanted cohort, as was done for the acute cohort. The cutoffs are indicated for each animal for the lowest (Fig. 2, solid vertical lines) and highest (Fig. 2, dotted vertical lines) SPLs recorded. In vivo (blue) and postmortem (black) cutoffs were estimated. The mean −3 dB cutoff re: peak displacement value for in vivo recordings at threshold was 264.02 Hz ± 78.004 Hz (N=12). At 40 dB SPL above threshold, the mean value was 455.78 Hz ± 76.58 Hz (N=5). Postmortem, the mean low SPL cutoffs were 612.24 Hz ±188.11 Hz (N=12) and 399.73±35.76 Hz (N=5) at high SPL. There was no statistical difference between the −3 dB cutoff frequencies measured at response threshold between the acute and chronic cohorts (acute pre-CI vs. chronic CI, p=0.6786, df=14; acute CI vs. chronic CI, p=0.2537, df=14, Student’s 2 tailed T test).

#### Apical organ of Corti sensitivity and non-linearity (Chronic cohort)

The contribution and condition of the cochlear amplifier is simpler to assess if it is expressed in terms of its gain relative to the middle ear measurement reference (in this case, the ear canal pressure). Figure 2 shows the iso-intensity sensitivity curves relative to ear canal pressure from 12 preparations where both in vivo (blue) and postmortem data (black) was collected. Broadly, there are two overall outcomes after chronic implantation. In 4/12 animals, tone complex evoked organ of Corti mechanical response thresholds were as low as those of the acutely implanted cohort (Fig 2. A, B, K, L). As such, data were collected only up to 74 dB SPL. The remaining preparations (Fig. 2 C-J), displayed response thresholds 30-50 dB higher than the lowest detected threshold (34 dB SPL). In these cases, data was collected for higher SPLs, although the linearity of the speaker system likely diminishes above 74 dB SPL. All preparations indicated at least some gain relative to postmortem responses, and 2/5 of the preparations with increased thresholds (Fig. 2 C, J) displayed physiologically vulnerable (therefore of biological origin and not attributable to sound system non-linearity) compressive non-linearity.

#### Cluster analysis of the apical organ of Corti sensitivity

Figure 3 shows that the residual hearing sensitivity for the chronic cohort varied. To identify the source of this variability, we performed k-means cluster analysis on the acute-post CI (red solid lines) and chronic organ of Corti (blue solid lines) sensitivities at the lowest SPL indicated in Fig. 1 & 2, with acute pre-CI (black solid lines) included for visual reference but excluded from clustering analysis. Fig. 4A shows that we identified two clusters (Fig. 4B, red and blue dots, centroids indicated by red and blue dashed lines. Calinski-Harabasz index = 57.501). The top cluster compares impeccably well with the same recordings made in (28) when accounting for the boosted low-frequency sensitivity derived from the closed cochlear condition here.

**Figure 3:**
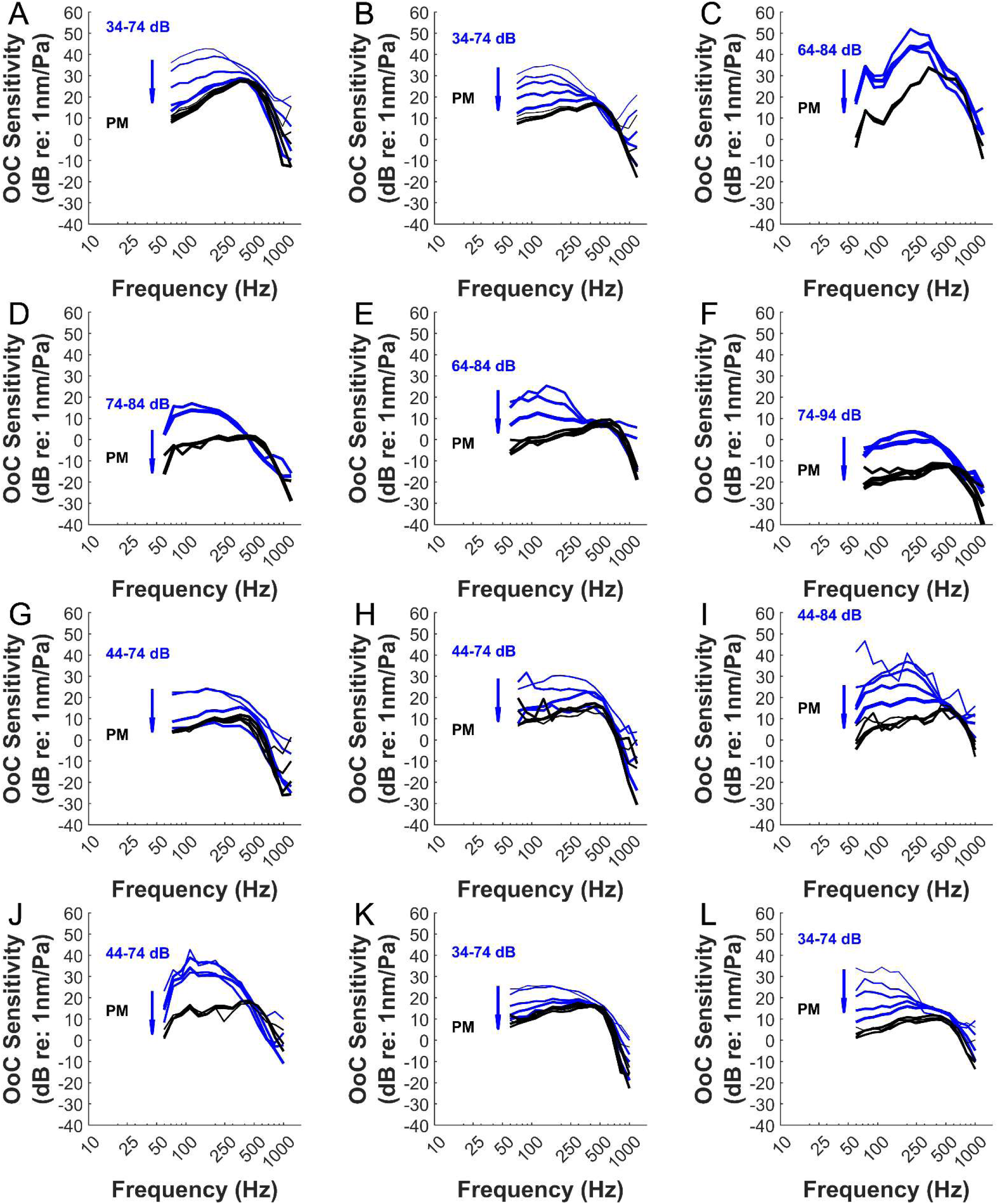
Iso-intensity sensitivity functions relative to ear canal pressure, in the 12 chronically implanted animals (A-L) shown in figure 2, in vivo (blue lines) and after euthanasia (black lines). Blue arrows indicate direction of gain change as SPL increases.

**Figure 4:**
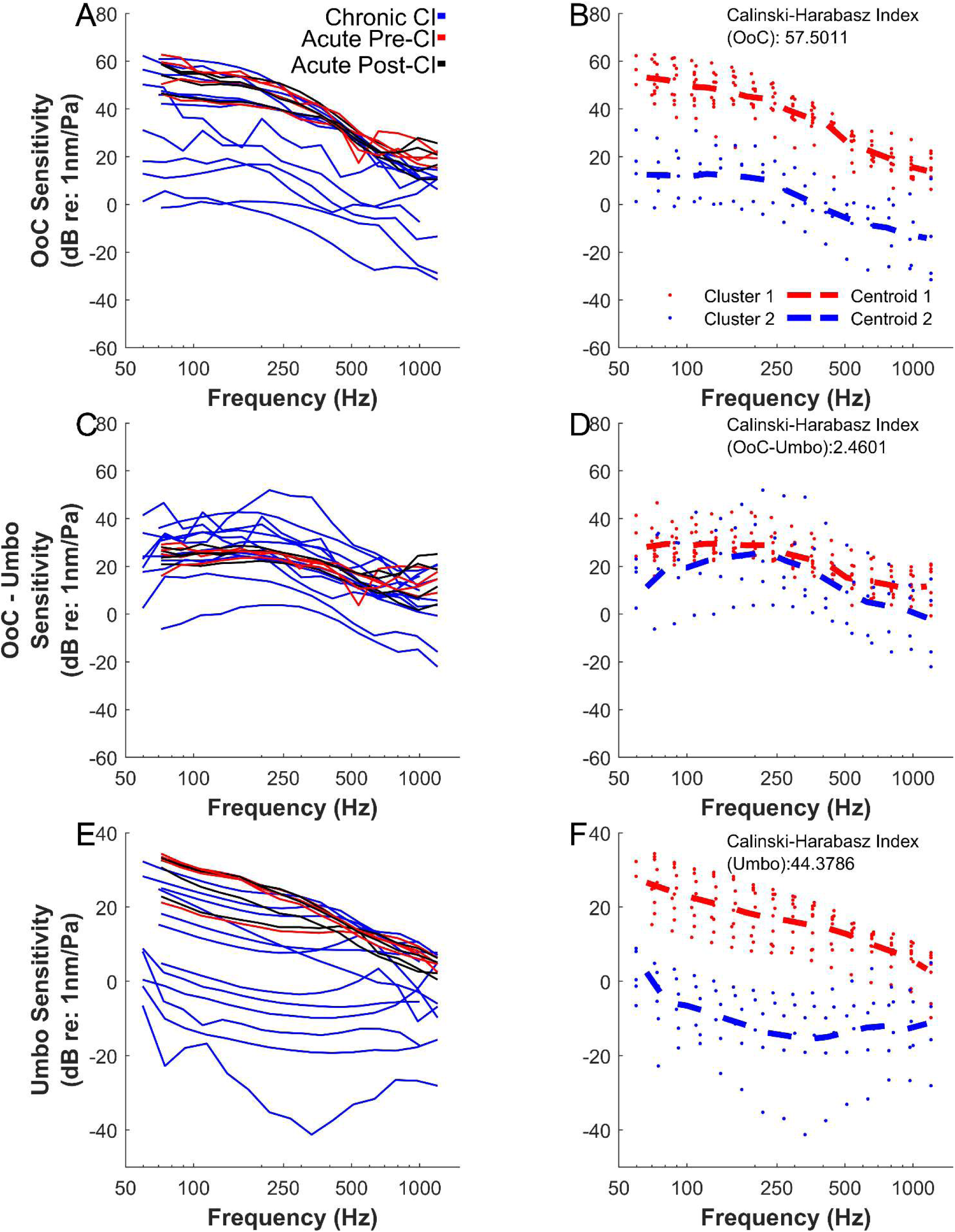
A: Organ of Corti iso-intensity sensitivity function re: ear canal pressure, for chronically implanted animals (blue), acutely implanted animals pre (red) and post (black) CI. B: A k-means cluster analysis revealed two clusters (red and blue circles, dashed red and blue lines = centroids). A Calinski-Harabasz index value of 57.501 indicates strong clustering. C: As in A, but subtracting the sensitivity of the Umbo. D: The same members of each cluster are identified as in A. Now, the Calinski-Harabasz index is 2.460, indicating clustering collapse. E. Umbo iso-intensity sensitivity function re: ear canal pressure (colors as in A and C). F: Cluster analysis via k-means revealed two clusters, with a Calinski-Harabasz index of 44.380, indicating strong clustering.

As we remarked above, gain is present despite increased threshold. Hence, we hypothesized that summing the organ of Corti and Umbo sensitivities would lead to declustering, identifying the middle ear as the main source for the bimodal distribution of sensitivities. This outcome would point to the presence of conductive hearing loss. Fig. 4C shows the sensitivity of the Organ of Corti minus that of the Umbo for each preparation. Fig. 4D shows the same cluster identities calculated in Fig. 4B, with umbo-corrected values and centroids. These two clusters are no longer separated (Calinski-Harabasz Index = 2.460, where a lower relative value for the same cluster populations indicates worse clustering). Hence, the umbo sensitivity variability explains most of the differences seen in the chronic cohort apical mechanics.

#### Umbo Sensitivities (Chronic cohort)

Fig. 4E shows the umbo sensitivities relative to ear canal pressure for each cohort, colored as in Fig. 4A-B (Fig. S1 shows the sensitivities arranged as in Fig. 2, 3 and 5). The acutely implanted animals had the largest sensitivities which changed little from pre-(red) to post (black) CI. Chronically implanted umbo sensitives varied and appeared to cluster. Fig. 4F shows a k-means cluster analysis identified two strong clusters (Calinski-Harabasz Index = 44.3786). Again, comparisons between the upper, red cluster in Fig. 4E-F and those of (28) are encouraging, save that overall sensitivity should be lower here as it is measured earlier in the ossicular chain. As we concluded above that most of the residual hearing loss in the chronic cohort is caused by conductive effects, we will assume that cluster 2 (green circles) represents the sub-population with residual hearing loss for further analysis.

**Figure 5:**
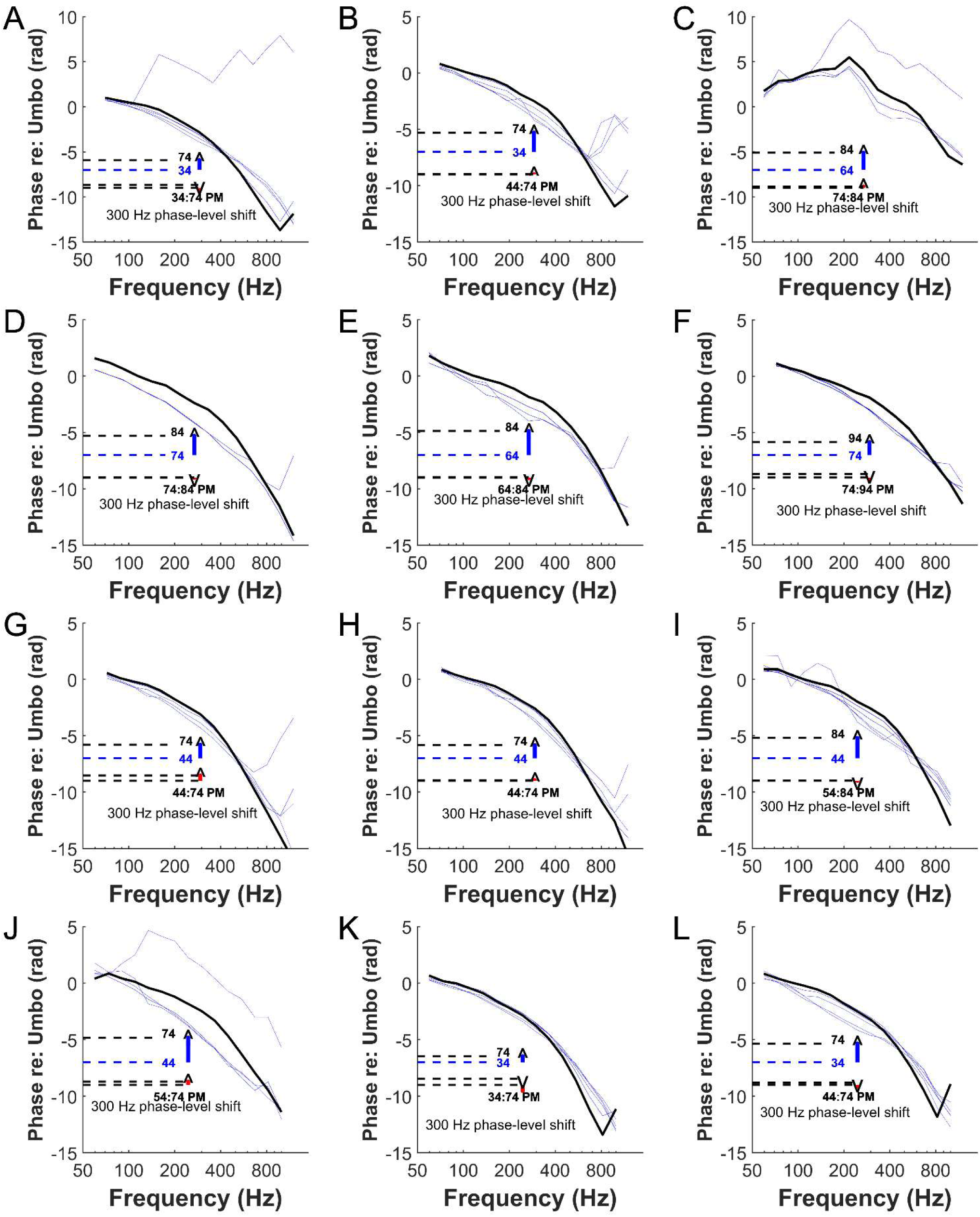
Organ of Corti phase referred to the umbo phase for each chronically implanted animal in figures 2-4 (A-L). Phase is shown in vivo (blue) and post-mortem (black) responses. As the SPL range used is not comparable, the level dependent phase shift is indicated using the difference between in vivo low SPL (dotted blue line) and postmortem (dashed black lines) phase. Postmortem phase at 300 Hz is indicated and is less variable with SPL (black dashed lines).

#### Phase of the organ of Corti response (Chronic cohort)

Figure 5 shows the phase of the organ of Corti response referred to that of the umbo for each experiment shown in Figs. 2-4. The phase is shown for the in vivo response (Fig. 5, blue curves) and the postmortem response (Fig. 5, black curves). As with the acute cohort (Fig. 1), there is evidence of a progressive phase lead with increasing SPL, indicated by the offset dashed lines. However, since the full dynamic range of the organ of Corti could not always be collected, we calculated the full magnitude of the phase shift as the difference between the in vivo lowest SPL response (blue horizontal dashed line) and the post mortem response (black horizontal dashed lines), which is not level dependent.

Importantly, the increased threshold of some of the experiments (E.g. Fig. 5D, F) does not lead to a diminished phase shift as might be expected from diminished cochlear amplification. Hence, the phase responses support the notion that chronic cochlear implantation has primarily induced conductive hearing loss in these experiments.

In summary, the apical organ of Corti mechanical response evoked from chronically implanted guinea pigs indicates a functional cochlear amplifier whose sensitivity and dynamic range is primarily limited by conductive mechanisms, where residual hearing loss is present.

### Statistical analysis of the mechanical indices of residual hearing and its loss

The key features highlighted above (response threshold, the gain of the cochlear amplifier, raw displacement at the organ of Corti and middle ear) are compared using correlation analysis in Figure 6. The relationship between organ of Corti gain

**Figure 6:**
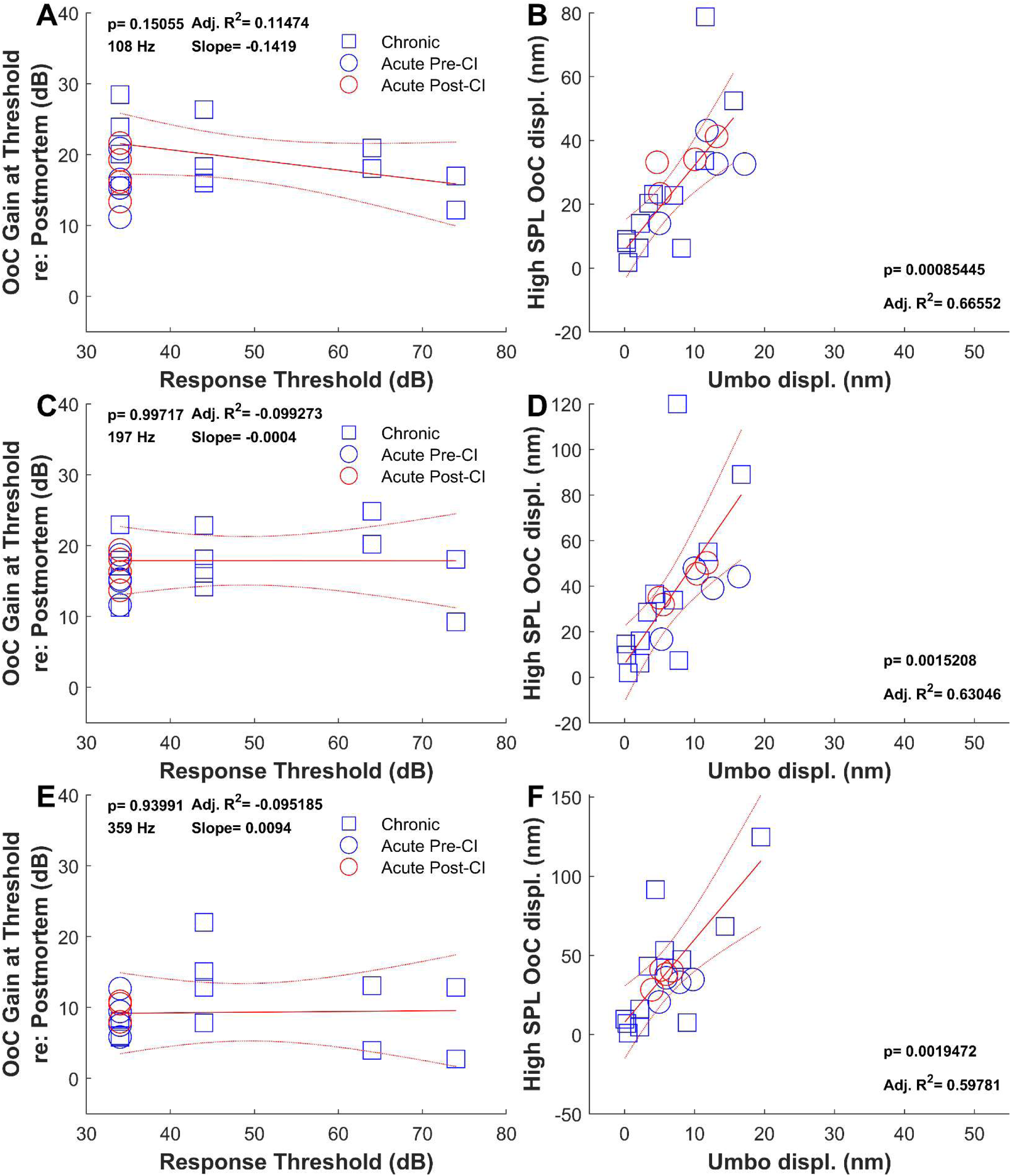
A, C, E: Gain value of the in vivo organ of Corti re: postmortem values, as a function of the response threshold of the organ of Corti for 104 (A), 192 (C) and 355 Hz (E) respectively, fitted by a linear regression (black line, curved dotted lines showing 95% confidence intervals) of the chronically implanted cohort (Squares). Acute pre-(blue circles) and post (red circles) CI gain values are shown for reference. B, D, F: Organ of Corti displacement as a function of the equivalent umbo displacement for the highest recorded SPL for each preparation, for 104 (B), 192 (D) and 355 Hz (F) respectively, fitted by a linear regression (black line, curved dotted lines showing 95% confidence intervals) of the chronically implanted cohort (squares). Acute pre-(blue circles) and post (red circles) CI gain values are shown for reference.

(Fig. 6 A, C, E, relative to postmortem displacement, which takes into account differing umbo displacements) and response threshold were compared at 104 (Fig. 6A), 192Hz (Fig. 6C), and 355 Hz (Fig. 6E). These frequencies were selected to exhibit low, medium and high frequency (within the apical turn) organ of Corti behavior which display differing non-linearity and amplification. The amount of gain produced by the cochlear amplifier in chronically implanted animals (blue squares) is not significantly correlated to the lowest displacement threshold at any of these three frequencies (Fig. 6A: 104 Hz, X/Y=-0.14 Adj. R^2^=0.114, p=0.15, C: 192 Hz, X/Y=0.0004, Adj. R^2^=-0.0992, p=0.997, E: 392 Hz, X/Y= 0.0094, Adj. R^2^=-0.0951, p=0.940). The pre- and post-CI gain values (Fig. 6 A, C, E, blue and red circles respectively) calculated for the acute cohort fall along the regression line, but there is no spread in threshold. Hence, the acutely and chronically implanted animals share similar gain, there is significant spread in the threshold of the organ of Corti displacement in the chronic cohort, but increased threshold does not correlate with reduced gain, as would be expected if there was a sensory component to the residual hearing loss at the apex.

Increased threshold in the presence of intact organ of Corti amplification strongly suggests that the residual hearing loss exhibited by some of the chronic cohort is conductive in nature, i.e. that there is some change to either middle ear or inner ear access impedance, resulting in reduced energy of the travelling wave that propagates from the stapes to the apex relative to the input sound pressure level. To test this hypothesis, we assessed the correlation between the high SPL raw displacement of the apical organ of Corti, which minimizes the contribution of the cochlear amplifier due to passive mechanisms dominating, and the displacement of the umbo at the same SPL and at the same frequencies as in Fig. 6 A, C and E (104, 192 and 355 Hz Fig. 6 B, D, F respectively).

The umbo displacement is positively correlated with the organ of Corti displacement at all three of these frequencies in the chronic cohort (Fig. 6 B: 104 Hz, Adj. R^2^=0.67, p=0.00086, D: 192 Hz, Adj. R^2^=0.63, p=0.0015, F: 392 Hz, Adj. R^2^=0.598, p=0.002). Hence, the acoustic admittance of the umbo is a dominating factor in the prediction of the organ of Corti response. In summary, the lack of relationship between organ of Corti amplification and its displacement threshold coupled with a strong relationship between the unamplified displacement of the organ of Corti and the middle ear, indicate that the majority of the residual hearing loss in the chronic CI model is conductive in nature.

#### Effect of Electrical Stimulation

3 out of 12 of our chronic cohort experienced controlled chronic electroacoustic stimulation (Fig. 2, 3, 5 A-C). However, we often experienced electrical failures of the implant array, and therefore proceeded with inactivated implant placement for the remaining animals of the cohort. To assess whether the electrically stimulated animals are comparable to the rest of the cohort, we compared stimulation status to −3 dB re: peak frequency, level dependent phase change. Stimulation status did not cause a statistically significant change in any of these values (−3 dB re: peak frequency: p=0.917, F=0.011, Linear Mixed Model; Level dependent phase change: p=0.2661, F=1.25, Linear Mixed Model). Also, we checked whether electrical stimulation was associated with changes in response threshold or gain (As correlated in Fig. 6). We did not find a statistical difference between the stimulated and unstimulated groups for either threshold or gain (p=0.549 and p=0.6590 respectively, Student’s 2-tailed T-test). Therefore, we chose to include all 12 of the chronic cohort in our analysis.

### Comparison of organ of Corti mechanical behavior between acute, chronic and residual hearing loss subjects

The organ of Corti mechanical response thresholds varied greatly between chronically implanted animals (Fig. 2). Additionally, cutoff values appear to increase as SPL increases in both the acute (Fig. 1) and chronic (Fig. 2) groups. However, since the apparent dominant contributor to this threshold spread was the evoked displacement at the umbo (Fig. 4) and was no relationship between threshold and the calculated gain values (Fig. 6), we could now compare the mechanical tuning based upon the assumption that the mechanical response characteristics at threshold are comparable. Further, to establish whether any animal statistically exhibited residual hearing loss, outlier detection using Median Absolute Deviation method was conducted upon the acute and chronic cohort thresholds. 4/6 animals from the chronic cohort were identified as possessing mechanical thresholds that were statistical outliers from the group (Fig 2. C-F). All of these animals were included in Cluster 2 (Fig. 4). Only 1/6 Cluster 2 animals received electrical stimulation. For later analyses, the animals in Cluster 2 are considered to have residual hearing loss.

Based upon these assumptions, we determined whether chronic placement of the CI leads to changes in local tuning characteristics in the cochlear apex by comparing −3 dB cutoff re: peak displacement values between the chronically implanted group without residual hearing loss, the pre-CI acute group and those among the chronically implanted group deemed to exhibit residual hearing loss. We constructed a linear mixed model (Cutoff_Frequency ∼ SPL_Above_Threshold x Implant_Status + RHL + Stim + (1 | Subject)). Figure 7 shows the distribution of −3 dB re: peak displacement of the organ of Corti versus the input SPL of that tuning curve. There was a statistically significant relationship between SPL and cutoff frequency for the chronic cohort (Fig. 7, blue circles; SPL versus cutoff: p= 8.71×10^-8^, F=37.25, LMM).

**Figure 7:**
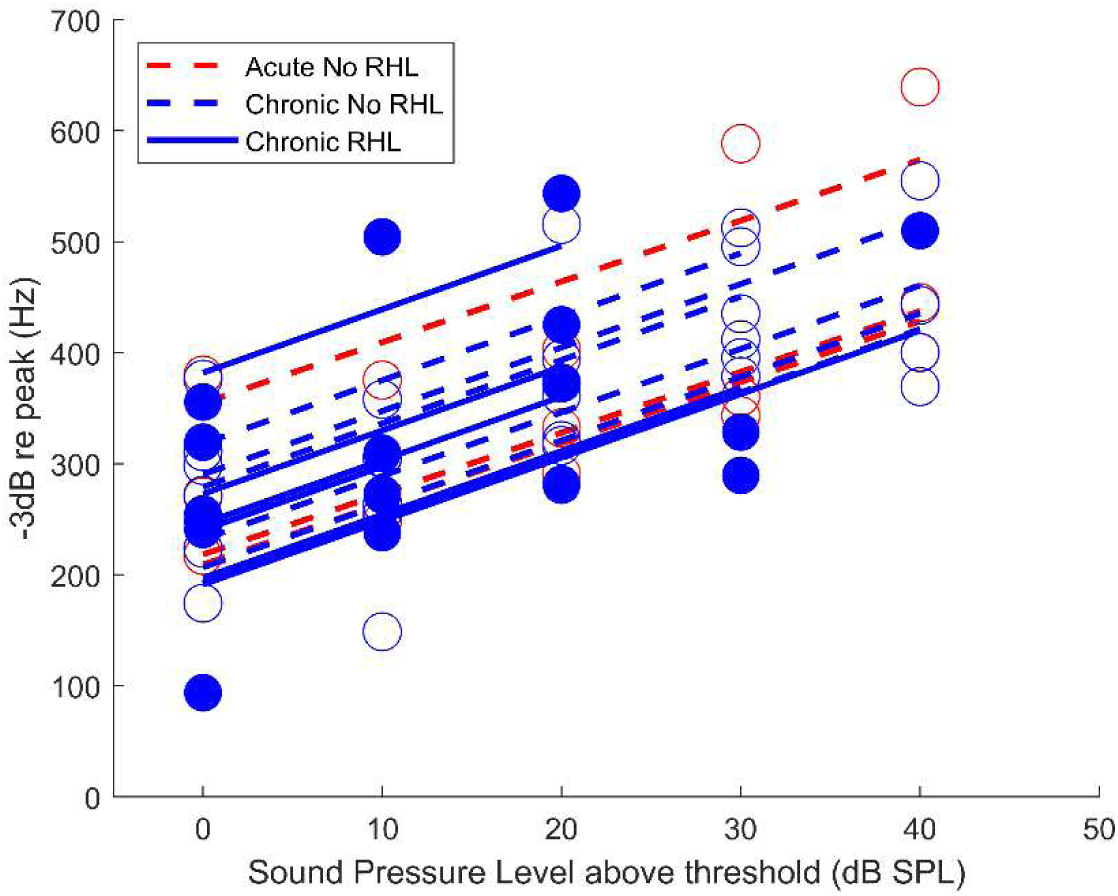
-3 dB re max displacement as a function of sound pressure level for the acute pre-CI (red) and chronically implanted (blue) cohorts, fitted by a generalized linear mixed model (red and blue lines respectively). Solid lines indicate animals determined to have residual hearing loss via the median absolute deviation outlier detection method.

However, implantation status did not impact tuning change with SPL (SPL vs. implantation status: p=0.895, F=0.0176, LMM), meaning that the level dependence of tuning did not depend on whether an implant was chronically present. Finally, the presence of residual hearing loss did not statistically impact tuning or its change with SPL (p=0.963, F=0.002, LMM). Hence, chronic CI had no influence on the local mechanical tuning properties of the apical organ of Corti in animals with or without residual hearing loss.

As the phase of the organ of Corti response was established to be a signifier of cochlear amplifier integrity in the cochlear apex (Fig. 1, 5), we fitted the level dependence of the phase data using a linear mixed model. Figure 8 shows the phase versus SPL above threshold for the chronic in vivo (blue circles), chronic postmortem (yellow circles), acute pre-CI (gray circles), Acute CI (Red circles) and Acute postmortem (black circles). Lines of corresponding color indicate the fits of the LMM formulated thus: (Phase re: Umbo ∼ 1 + SPL x Implantation_Status + RHL + Stim + (1 | Subject)), where Implantation_Status can refer to each of the 5 conditions listed above, and where RHL is a covariate that denotes whether the mechanics indicated residual hearing loss (solid lines) or not (dashed lines) as described above.

**Figure 8:**
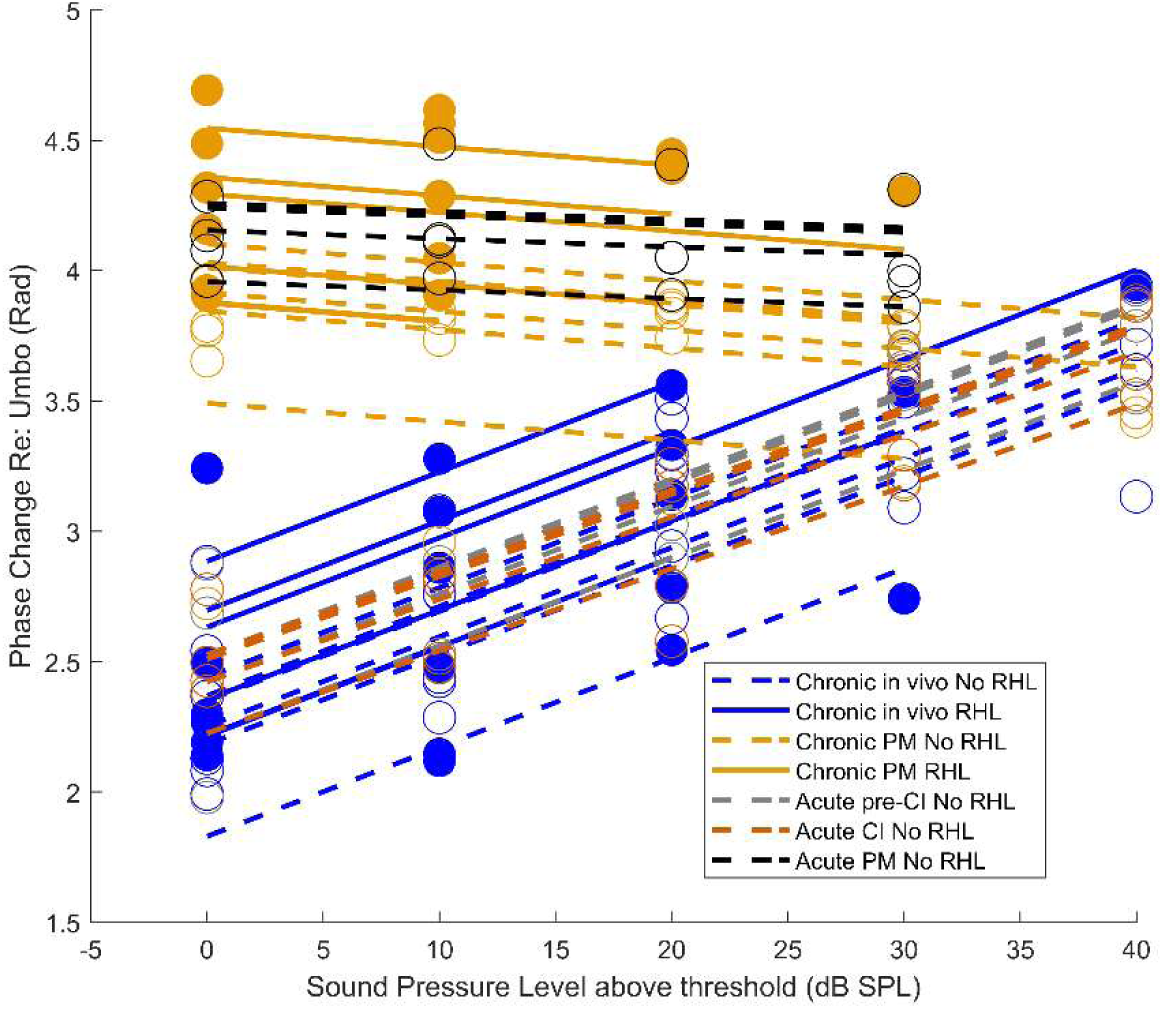
Relationship between SPL above threshold and organ of Corti phase re: umbo for Chronically implanted animals in vivo (blue), Chronically implanted animals postmortem (yellow), Acutely implanted animals pre-CI (gray), Acutely implanted animals post-CI (red), and Acutely implanted animals postmortem (black). Dots are datapoints, and lines are individual fitted slopes based upon the output of a linear mixed model. Solid plots indicate animals where residual hearing loss was identified using the median absolute deviation outlier detection method.

Euthanasia led to a strongly significant decrease in the slope (Chronic Postmortem: p=1.323×10^-18^; Acute Postmortem: p=3.920×10^-18^, LMM) and intercept (Chronic Postmortem: p=6.903×10^-45^; Acute Postmortem: p=5.791×10^-19^, LMM) of the phase-SPL relationship. However, in vivo pre-CI nor acute CI conditions did not lead to significant changes in that slope (acute pre-CI: p=904; acute CI: p=0.560, LMM) or intercept (acute pre-CI: p=0.314; acute CI: p=0.310, LMM). The presence or absence of RHL was not a statistical predictor for changes in phase-SPL (p=0.0598, F=3.602, LMM).

A coefficient test of the slopes produced by the model confirmed that the in vivo and postmortem phase-level function did not correlate in the chronic dataset (p=4.750×10^-40^, F=377.28, linear coefficient test with Bonferroni correction), as expected given the diminished phase lead with SPL following euthanasia (Fig. 5). In contrast, the chronic in vivo group did not differ significantly from either the acute pre-CI or acute CI groups (vs. acute pre-CI: p=1, F=1.2025; vs. acute CI: p=1, F=0.782, linear coefficient test with Bonferroni correction). Further, there was no significant difference between acute pre-CI and acute CI slopes, (p=1, F=0.190, linear coefficient test with Bonferroni correction). These coefficient test results suggest that the local non-linear characteristics of the cochlear apex are unaffected by acute or chronic CI.

The statistical analyses above demonstrate that the acute or chronic presence of a CI, or identifying residual hearing loss by umbo sensitivity (and to a lesser extent to threshold outlier status) does not isolate changes in the mechanical responses of the apical organ of Corti that are tied to its local, physiologically vulnerable activity.

## Discussion

This study provides the first successful mechanical recordings of residual hearing from the intact, living cochlear apex of an acute and chronic cochlear implantation model. We concluded that the presence of the CI neither acutely nor chronically impacts the local function of the cochlear apex in guinea pigs, since CI placement acutely had minor effects upon the apical cochlear amplifier (Fig. 1), there was no statistical relationship between the organ of Corti gain and displacement threshold (Fig. 5 A, C, E), and there was no statistically significant change in the relationship between SPL and −3 dB re: peak displacement caused by the chronic presence of an implant (Fig. 6). However, the sensitivity of the middle ear is responsible for the separation of the OoC responses into lower and higher sensitivity clusters (Fig. 4). Also, there is a significant correlation between the high SPL (passive dominated) displacement of the OoC among chronically implanted animals (Fig. 5 B, D, F). Hence, in this model, the dysfunction of the cochlear apex was primarily conductive in nature. Conductive changes to patient residual hearing post CI support the clinical relevance of this finding (35, 36), especially since we may directly observe that conductive loss’ impact upon organ of Corti function at the cochlear apex.

In a previous study using guinea pigs, 500 Hz cochlear place performance measured by auditory nerve overlapped waveform suggested that acute CI placement does not lead to conductive hearing loss at the cochlear apex. Reversible conductive hearing loss was detected at higher frequencies via compound action potential thresholds (20). It was noted that the spatial buildup of amplification was too restricted to impact apical amplification in the case of partial conductive loss – in other words, the placement of the CI did not impede the process of amplifying the travelling wave; a notion that our mechanical data from the apical turn can support. We only examined the apical cochlear place and corresponding response frequencies at the middle ear and cannot comment on high frequency changes. Our acute umbo data partially suggest that conductive hearing loss can result from acute CI placement (some small decreases were noticed e.g. Fig. 1L), but this may be a peculiarity of rodents rather than humans for two reasons. First, the guinea pig scala tympani area taken up by the CI is significantly larger than in humans (22), and in humans the CI electrode will not fill the space, reducing the likelihood of direct contact with the organ of Corti. Second, stapes/round window measurements made perisurgically prior to and following CI did not reveal any systematic, instantaneous change in displacement that could not be attributed to measurement error (37). Preserved non-linearity at the cochlear apex would point to hydromechanical changes induced by the presence of the CI.

CI placement in humans is associated with pressures equivalent to extreme SPLs (38–41), and hence our observations of loss of non-linearity or gain in the acute cohort are entirely consistent with surgical trauma. A limitation in the acute experiment is that we did not withdraw the CI to examine reversibility of the mechanical changes. Hence in future experiments, it would be ideal to also record the influence of array removal upon apical and middle ear mechanics.

Approximately half of the chronically implanted cohort displayed significant residual hearing loss, indicated by increased response threshold in the cochlear apex and at the middle ear (and hence reduced displacement at higher SPL). At no point did we observe cochlear dysfunction without accompanying middle ear dysfunction in the chronic cohort. Note that it is possible that the impact of the observations made here may be underestimated in humans, because the guinea pig cochlear implant used (8 mm) was significantly shorter than the ∼20 mm long guinea pig cochlear duct (22). In the human cochlea, there is substantially more overlap between the implant and the organ of Corti length. Hence, the human case, where there is more apical electrode presence and ensuing fibrosis, the spatial extent of the changes in mechanical environment caused by fibrosis and array may further influence travelling wave propagation.

Pathohistology of the organ of Corti was not explored in this study, as the substantial postmortem measurement period was likely to impact the quality of tissue for histology. The extent and distribution of the intracochlear fibrosis in these cochleae was recorded, and it is the subject of ongoing analysis for a future study. However, previous studies on guinea pigs indicate that outer hair cell loss is not a driver of residual hearing loss (14, 15), especially in younger animals (24). Our apical mechanical findings support this notion as non-linearity and amplification was intact in the apical turn of almost all chronically implanted animals.

This study therefore indicates that significant conductive dysfunction can contribute to residual hearing loss, and that basal turn electrode presence can lead to that conductive dysfunction. One may interpret the reduced displacement of the umbo in response to a given SPL due to implant presence is a result of increased middle ear impedance. That increase in impedance may be due to the increased cochlear access impedance, or by changes in middle ear function (e.g. otosclerosis). A limitation of this study is that otosclerosis was not systematically investigated, although there was no impression that this was common. Also, it was not possible to measure the function of the stapes as the bone was obscured by the CI, and the wider dissection necessary to expose it may disturb the array and lead to additional confounding trauma. Future studies would benefit from audiometric measurements of middle ear function, such as wideband emittance or tympanometry. Additionally, the source of the impedance changes seen in this study may be explored experimentally in future OCTV measurements. For example, we suggest measurement of organ of Corti and middle ear displacement before and after the placement of a third window in the scala tympani, apical of the implant location. This third window should induce a drop in cochlear mechanical access impedance if the source is intracochlear, and not if the source is extracochlear e.g. from middle ear issues.

An unfortunate limitation we encountered was technical difficulties in our chronic electrical stimulation protocol. We found that, sometimes even prior to stimulation, electrode arrays could cease to function. Given that the goal of this study was to examine the mechanical consequences of foreign body response, we elected to continue with inert electrode arrays. The statistical tests detailed above, while limited in power by small N, did not identify differences caused by electrical stimulation, and so we chose to include the stimulated animals in our analysis. A future study with a larger electrical stimulation arm will be of great use in examining the mechanical consequences of electrical stimulation. However, similar guinea pig studies using conventional physiological techniques found no statistically significant evidence of apical hair cell dysfunction associated with electroacoustic stimulation (15, 16). Notably the latter study pointed to the need to assessment of conductive hearing loss; something we have prioritized in the current study.

In summary, we have given guinea pigs acute and chronic CIs, and successfully recorded non-linear, amplified organ of Corti responses at their cochlear apexes, as well as responses from the umbo of the tympanic membrane. This experiment provides direct information upon the function of the implanted cochlear amplifier, and could be used as a testbed for objective, direct appraisal of potential therapies for the treatment/prevention of residual hearing loss, or for testing the mechanical impact of novel implant types. Our conclusions from this study support the idea that cochlear access impedance increases following CI, and may cause conductive hearing loss (6). These findings encourage alternative acoustic stimulation strategies such as bone conduction EAS, and also point to mitigation of mechanical changes being an effective means to mitigate residual hearing loss following CI, in concert with other treatments.

## Acknowledgements

The authors would like to thank Dr. Anders Fridberger for the use of his OCTV acquisition and analysis software, Dr. Bradley Buran for his advice on statistical modeling, Dr. Tianying Ren and Dr. Teresa Wilson for their helpful comments on earlier versions of this manuscript. The authors also thank Cochlear inc. for provision of purchased implant arrays and manuscript feedback. This work was supported by grants from the National Institute for Deafness and Communication Disorders (R21DC020794-03, Burwood; R56DC018387, Reiss), and the Hearing Health Foundation (ERG969672, Burwood).

## Methods

### Animals

All experiments were conducted in accordance with OHSU IACUC policies and with the permission of the local ethics committee (Protocol numbers IP00001278 & IS00000672). 25 male albino guinea pigs were used in this study (20 chronically implanted and 5 acutely implanted), and data from 16 are included in this paper. 2 guinea pigs did not survive CI surgery due to peri-surgical complications. 1 guinea pig was euthanized during its post-operative monitoring following CI surgery due to poor health outcomes necessitating a humane endpoint. 1 guinea pig was euthanized 4 weeks after CI surgery due to meeting the criteria for a humane endpoint. Full in vivo and postmortem datasets were not successfully collected from a further 3 animals during OCTV measurements, and 1 further animal was found to have experienced explantation at an unknown timepoint upon opening of the bulla for OCTV experiments. One acutely implanted guinea pig was excluded from the dataset as implantation led to damage of the tympanum.

### Cochlear implant surgery

The surgical approach was previously described in (24). Briefly, animals were given water to wash hay or food from cheek pouches, and pre-treated with analgesia (either Buprenorphine or Meloxicam) initially with ketamine (30 mg/kg IM) and xylazine (3-5 mg/kg IM). The surgical field was shaved and the animal was placed upon a servo-controlled heating blanket with a rectal temperature probe placed. Oxygen and isoflurane (dosing: 2%-3% in 100% oxygen with a flow rate of 1.5-3L/min; increase to 3% during incisions; decreased to 2%-2.5% after major surgical procedures such as drilling, and dissecting are completed; decreased to 1.5%-2% during suturing; flushed with oxygen for a few minutes after completion of surgery) was provided to maintain areflexia for the duration of the procedure. Heart rate, spO2 and breathing rate were monitored using a Cardell veterinary monitor. A single surgeon performed implantation surgeries (WH). Using aseptic technique and under a Zeiss operating microscope, the postauricular skin was incised and muscle was blunt dissected to reveal the posterior temporal bulla. This was opened with a diamond burr and the round window was visualized. A diamond burr was used to place a cochleostomy at the edge of but not contiguous with the round window membrane. An HL08 pre-clinical cochlear implant array (Cochlear Ltd., Sydney, Australia) was slowly introduced into the cochleostomy until resistance was met. Excess implant lead wires were trimmed and coiled into the posterior bulla, away from the middle ear. A muscle plug was placed over the bullostomy and the incision was closed with absorbable sutures. Anased was administered to reverse xylazine, and the animal was maintained on low isoflurane while its nails were trimmed and it was administered warmed lactated Ringer’s solution. Finally, the isoflurane was switched off, and the animal was maintained on the heat pad on oxygen support until it showed signs of waking. At this point, the animal was wrapped in a blanket and transferred to a warm, clean cage to recover from anesthesia.

Where electrical stimulation was received, the surgical procedure differed – a serial port was fitted to the top of the skull with screws, and signal and ground wires were guided under tissue to the posterior bullostomy to provide electrical stimulation to the array. For more details, see (16).

The postoperative monitoring procedure typically lasted 3 days, where the animal was given analgesia and enrofloxacin IM, and topical Silver Sulfadiazine antiseptic. The incision site was inspected for signs of infection, and the animal’s weight and body condition score were monitored. Animals were given Oxbow critical care diet supplement if necessary, and any animals that showed signs of infection, could not gain weight or did not thrive were referred for veterinary examination in consideration of reaching their humane endpoint.

### Electrical stimulation

Electrical stimulation was conducted as previously (16). Programming of the Freedom speech processor will be conducted using a programming pod, Freedom speech processor, emulator, and clinical Custom Sound software (Cochlear Inc., Sydney, Australia). The processors were programmed with monopolar stimulation using biphasic pulse trains, CIS processing strategy, 25 us pulse width, and 1800 pulses/sec. Frequency table 6 allocates the frequency range of 188-7938 Hz among the available electrodes. EABR thresholds were used to set T-levels, and C-levels were determined based on animal’s behavioral responses. The CI map was then written to the speech processor.

Due to a high incidence of implant electrode failures and other logistical challenges, we elected to cease electrical stimulation of the cohort. The three animals that received electrical stimulation were housed in tall, open cages that allow a tether cable system with slip-ring connectors to be mounted and connected to the percutaneous connector mounted on top of the animal’s head. The cable is counterbalanced, adjusted for weight and sprung to dampen movements. Electric stimulation was delivered via a direct audio input connection to a Freedom speech processor, which was connected to an implant emulator to convert the signal into biphasic current pulses. The current was delivered via the tether cable. The stimulus was amplitude modulated white noise (50% modulation, 65 dBA, 188-7938 Hz) stimulating a broad frequency range and dynamic, like speech. Animals were acclimated to the stimulation starting with 1 hour the first day, and doubling each day, until 8 hours was reached. Animals received 26, 29 and 29 days of stimulation respectively.

### OCTV surgery and measurements

We employed our minimally invasive surgery for exposure of the cochlear apex to measure the organ of Corti mechanical response using OCTV (25, 27, 30, 31, 33). Briefly, animals were anesthetized with ketamine and xylazine and were wrapped in a heating blanket. The temporal bulla was exposed via the ventral aspect and the anterior bulla was opened slowly and carefully using rongeurs. The posterior bulla region was left untouched as opening it was considered a risk to the integrity of the CI array. Cochlear surface vessels were removed using a saline-damped cotton pleget and a custom cut 2mm^2^ gold sputter coated cover slip square was placed on a 3D printed scaffold and placed at the apex of the cochlea, in order to direct the beam of the OCT microscope axially along the cochlea, permitting transverse measurement of the organ of Corti displacement. A Thorlabs Telesto III OCT microscope with a 5x objective lens (LSM03, Thorlabs Gmbh) was moved into position above the animal and the mirror was located using the OCT microscope’s camera. An OCT image of the cochlear apex was resolved by focusing the lens and the reference path and a scan line was drawn across the apex as visualized in the mirror. Displacement was measured in response to logarithmically spaced tone complexes between 60-1000 Hz and each frequency component at 34-94 dB SPL, with random phase, calibrated using an in-situ sound system. Vibration measurements were usually made by scanning the cochlea across the mid-modiolar plane, with the equivalent of 70 A lines per B scan across ∼2 mm, excluding irrelevant pixels (corresponding to the modiolus, for example). Measurements of the mechanical response to the stimulus were made until there was no detectable response visible from on-line inspection of the acquired spectra. Data were acquired from similar locations to those reported in (27), while the reported data are those with the most gain and non-linearity, corresponding to approximately 20% of the distance from the helicotrema. Umbo displacement data was acquired only for one SPL, as the middle ear response is linear.

For chronically implanted animals, the OCT microscope was put in place and not moved, but was refocused to resolve the umbo of the tympanum, which was visible in the same measurement plane as the mirror. For acutely implanted animals, baseline measurements were made and the OCT microscope was then moved to perform the cochleostomy and CI insertion, as above. Once the CI was implanted, the microscope was replaced using visual references through the CCD camera, and as close to the same measurement location as possible was reacquired. Following acute CI placement, the sound system was recalibrated.

Following completion of measurements, the animal was euthanized with an overdose of anesthetic. At least 30 minutes following euthanasia, measurements were repeated. Total experimental time could take between 6-8 hours.

### Data analysis

Data were analyzed using a modified approach and software developed for our mechanics studies (27). Briefly, the morphological image was used to identify the cellular portion of the organ of Corti. Displacement was calculated using algorithms developed in (42). All pixels whose mechanical response satisfied a 10 dB SNR were included in a database, and low-frequency noise was differentially subtracted using the in-line pixels corresponding to the cochlear bone. The top 10 most responsive pixels, usually corresponding to OHC-Hensen cell region, were used for analysis. An ROI was drawn on the umbo close to the surface of the bone, which has the strongest optical signal, and the top 10 most responsive pixels were used for analysis.

The median displacement of the pixels identified above were used to produce iso-level displacement tuning curves, and the *in vivo* or postmortem displacement values were divided by one another or by the ear canal pressure referred to 90mV/Pa (the 9 mV/Pa sensitivity of the in ear Knowles microphone with 20 dB of gain applied by its Etymotic ER-10 amplifier), to generate iso-level sensitivity tuning curves and local sensitivity estimates respectively.

To determine −3 dB high-side cutoff frequencies, in MATLAB, raw displacement tuning curves from the peak displacement value to the highest frequency value were fitted to a 4^th^ order polynomial function and interpolated, and the frequency corresponding to the −3 dB re: maximum displacement value was identified.

### Statistical analysis

Student’s 2 Tailed T Test, linear mixed models with post-hoc ANOVA, linear regression fitting and correlation testing adjusted for multiple comparisons were conducted. Significance was achieved at p≤0.05. All statistical testing was conducted in MATLAB. To determine whether mechanical response threshold had increased to outlier status and to quantitatively delineate animals with residual hearing loss, median absolute distribution test (Matlab: isoutlier) was used on the lowest SPL that evoked a recorded response in the organ of Corti, and k-means cluster analysis (where 2 clusters were visually assumed to be present) assessed with the Calinski-Harabasz Index, otherwise known as the variance ratio criterion.

---

**Figure S1:**
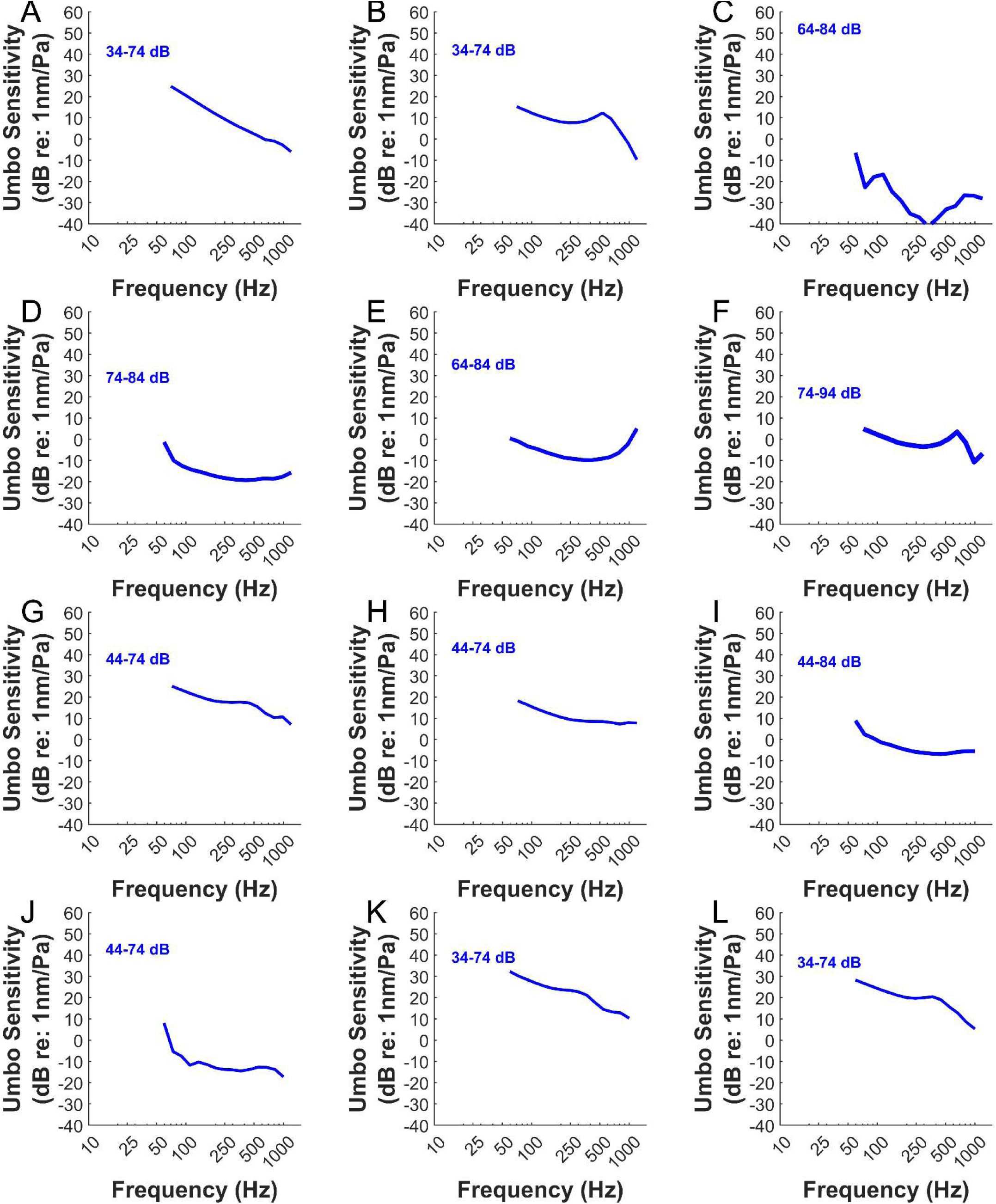
Iso-intensity umbo sensitivity functions relative to ear canal pressure, in the 12 chronically implanted animals (A-L) shown in figure 2, following euthanasia. The response range recorded from the organ of Corti is described.

